# Chemical effects on ecological interactions within a model-experiment loop

**DOI:** 10.1101/2022.05.24.493191

**Authors:** Dominique Lamonica, Sandrine Charles, Bernard Clément, Christelle Lopes

## Abstract

We propose in this paper a method to assess the effects of a contaminant on a micro-ecosystem, integrating the population dynamics and the interactions between species. For that, we developed a dynamic model to describe the functioning of a microcosm exposed to a contaminant and to discriminate direct and indirect effects. Then, we get back from modelling to experimentation in order to identify which of the collected data have really been necessary and sufficient to estimate model parameters in order to propose a more efficient experimental design for further investigations. We illustrated our approach using a 2-L laboratory microcosm involving three species (the duckweed *Lemna minor*, the microalgae *Pseudokirchneriella subcapitata* and the daphnids *Daphnia magna*) exposed to cadmium contamination. We modelled the dynamics of the three species and their interactions using a mechanistic model based on coupled ordinary differential equations. The main processes occurring in this three-species microcosm were thus formalized, including growth and settling of algae, growth of duckweeds, interspecific competition between algae and duckweeds, growth, survival and grazing of daphnids, as well as cadmium effects. We estimated model parameters by Bayesian inference, using simultaneously all the data issued from multiple laboratory experiments specifically conducted for this study. Cadmium concentrations ranged between 0 and 50 *μ*g.L^-1^ . For all parameters of our model, we obtained biologically realistic values and reasonable uncertainties. The cascade of cadmium effects, both direct and indirect, was identified. Critical effect concentrations were provided for the life history traits of each species. An example of experimental design adapted to this kind a microcosm was also proposed. This approach appears promising when studying contaminant effects on ecosystem functioning.

## Introduction

The toxic effects of contaminants are most often studied at the individual level, since it is easier to study life history traits of an isolated organism, studying for example its survival, development or capacity to reproduce. Moreover, monospecific bioassays are easy to implement and perform, and observation data at the individual level are straightforward to analyse, since they depict the direct effects of contaminants. Nevertheless effects can also be measured at other levels of biological organisation using various experimental devices that are chosen related to the level of interest, by adapting several characteristics such as size, duration, number of species, abiotic compartment, etc (Calow, 1993). Multispecies devices, like microcosms and mesocosms, allow to study organisation levels from populations to ecosystem, by integrating population dynamics and interactions between species (Forbes et al., 1997; Kimball and Levin, 1985; Ramade, 2002). However, extrapolating toxic effects from one biological level to the next based on observation data remains a challenge. In particular, going from individual to population levels, or from population to community levels, implies taking into account intra- and inter-specific interactions, which are of major importance in the functioning of ecosystems, while it is necessary to integrate these interactions for a better assessment of the ecotoxicological risk (Cairns, 1984; De Laender et al., 2008; Preston, 2002).

Modelling tools have proven their utility to analyse ecotoxicological data, by highlighting the underlying mechanisms leading to observations at each level of biological organisation. But modelling appears particularly helpful when extrapolation of contaminant effects from biological levels reveals necessary. For instance, physiologically based toxico-kinetic survival models allow to extrapolate the fate of a contaminant at sub-individual level to its effects on individual survival (Ashauer et al., 2016), or individual based models (IBM) including contaminant permit to extrapolate effects on the population level from effects on the individuals (Hansul et al., 2021; Mintram et al., 2018), or food web models permit to transfer effects of contaminants across the whole community (Baudrot et al., 2018).

Ecotoxicology relies on experimental data, while being concerned by the Replacement, Refinement and Reduction of Animals in Research (3Rs) program (Kilkenny et al., 2009) and by difficulties linked to collection of field data. Taking the most of experimental data and reducing the amount of experiments to perform in general is a key issue the ecotoxicology field faces. Formal optimisation of experimental design can be applied to standard tests (namely monospecific bioassays): they have been questioned in terms of test duration and measured endpoints (Charles et al., 2016) or regarding the tested concentration range (Forfait-Dubuc et al., 2012). Yet, more complex experimental designs, like microcosms or mesocosms may resist to formal optimisation particularly because of species interactions leading to indirect effects. Some attempts have been made in simple cases to deal with standard dose-responses curves (Chèvre and Brazzale, 2008; Holland-Letz and Kopp-Schneider, 2015; Keddig et al., 2015; Khinkis et al., 2003; Sitter and Torsney, 1995; Wang et al., 2006) but to our knowledge, nothing similar exist for multi-species models. Nevertheless, when modelling has been integrated to the experimental framework, it can easily be used to evaluate *a posteriori* the relevance of the data, as a pragmatic and case-by-case method to analyse the information provided by data and possibly improve the experimental design for studies with microcosm experiments with similar species and compounds.

The aim of our paper is to illustrate (1) how to use modelling to describe the functioning of a three-species microcosm exposed to a contaminant and to discriminate direct effects (related to effects on specific, modelled processes) and indirect effects (related to effects resulting from the cascade of processes); (2) how to develop critical effect concentrations for key population regulating processes (such as *EC*_50_ in stress functions); and (3) how model outcomes can inform experimental design in order to identify which of the collected data have really been necessary and sufficient to estimate model parameters in order to propose a more efficient experimental design for further investigations.

Different steps have been set up to achieve our objectives, as summarised in figure 1. We performed experiments to collect data on the microcosm species populations at different cadmium concentrations. In parallel, we formulated a model of the microcosm functioning under a chemical stressor based on coupled ordinary differential equations (ODE) and effect functions. First, using all data we estimated model parameters, in particular those related to effect functions (figure 1, black boxes). Using data where species occur in isolation and where they occur as a community of species permitted to identify direct and indirect effects of cadmium on the population dynamics of the different species. We then globally analysed the perturbations of our small community (objective 1). We also extracted *EC*_50_ for the different processes (growth, survival, and strength of interspecies interaction) (figure 1, orange boxes). In order to assess the relevance of certain data, we removed those data from the complete dataset to build partial datasets. Then, we estimated function parameters with the partial datasets. The newly estimated effect functions were then compared to the reference ones obtained with the complete dataset (figure 1, green boxes). This allowed us to evaluate the added values of only considering partial datasets instead of the complete original one (objective 2).

**Figure 1.**
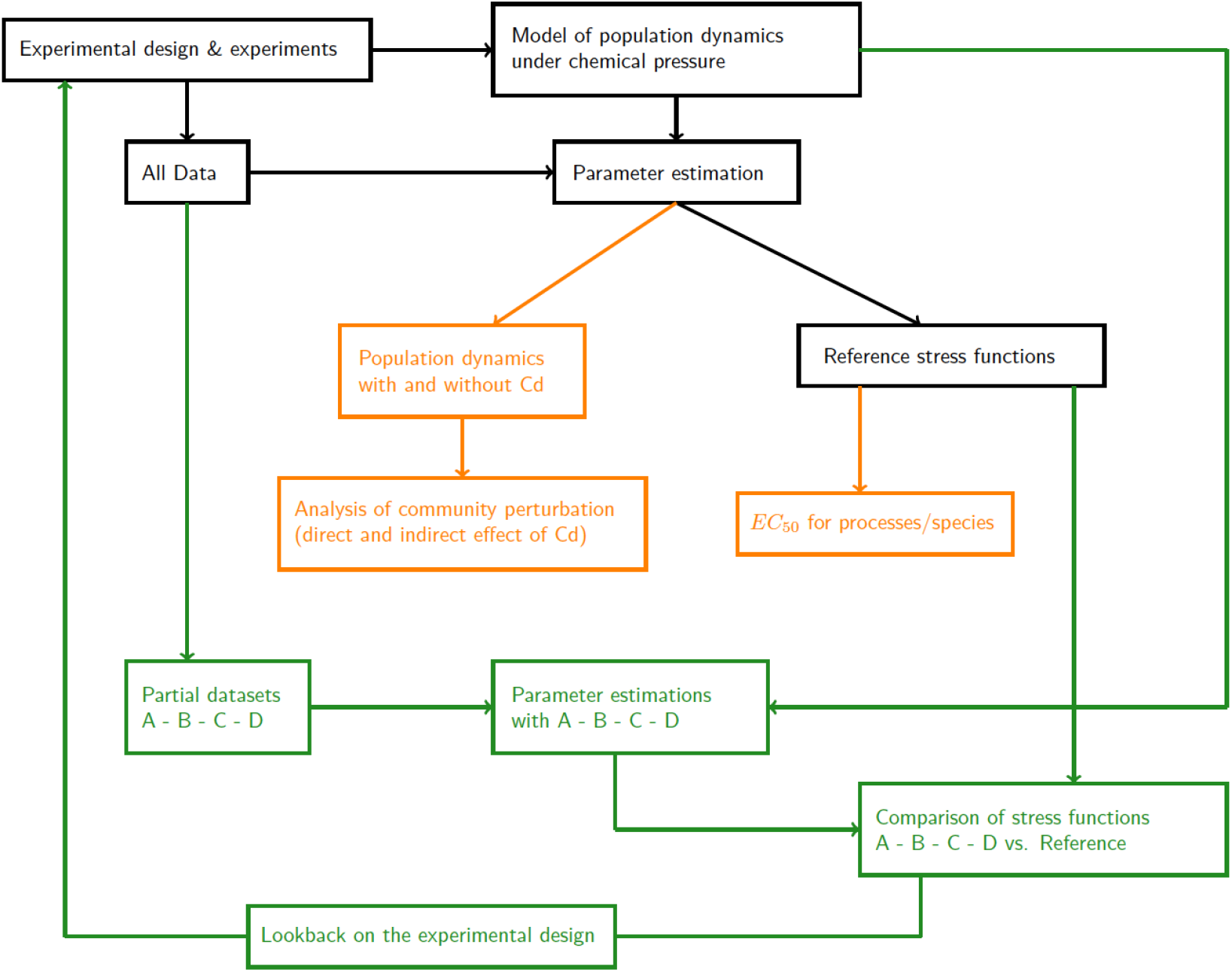
Workflow followed in the present study: (1) in black, development of the model of the microcosm functioning under cadmium pressure, data collection, and estimation of model parameters with the whole dataset, then extraction of stress functions, namely “reference stress functions”; (2) in orange, comparison of population dynamics with and without cadmium to analyse the global perturbation of the community, extraction of *EC*_50_ for the different processes; (3) in green, build of partial datasets by removing some parts of the complete dataset, estimation of parameters using those partial datasets, comparison of the newly estimated stress functions to the reference ones to assess how much information was provided by the removed data.

## Experiments and observed data

### Experimental design

Microcosms were identically prepared for all experiments according to (Lamonica, B Clément, et al., 2016) (without sediment). Algae, duckweeds and daphnids were cultivated at the laboratory according to internal protocols (BJP Clément et al., 2014). According to the experiment, beakers were inoculated with one, two or three species at the start of the experiment (day 0). When algae were present, 4.10^7^ cells of *P. subcapitata* were introduced into beakers. When daphnids were present, 10 daphnids (*Daphnia magna* neonates aged 24 *±* 12 h) were introduced into beakers. When duckweeds were present, 8 fronds of duckweeds were introduced into beakers. The algal density in the water column was measured every two to three days with a particle counter (Lamonica, B Clément, et al., 2016). The algal density at the bottom of the beakers was measured once during the experiment (Lamonica, Herbach, et al., 2016). Daphnids neonates were removed from the microcosm every two days, meaning that reproduction was considered as an independent process in the microcosm functioning (Lamonica, B Clément, et al., 2016). The number of daphnids in each beaker was counted (after neonate removal if necessary) and their size measured (from the centre of the eye to the caudal base of spine) twice or thrice per week. The duckweed fronds were counted every two to three days. The experiments lasted between 13 and 21 days. The experiments are summarised in SI Table S1 (Lamonica, Charles, et al., 2022c) and data can be found at (Lamonica, Charles, et al., 2022b).

### Experiments without cadmium

Experiment 1 involved algae and daphnids as detailed in (Lamonica, B Clément, et al., 2016) (referred in that paper as “Experiment without sediment” in section 2.3.2.). Experiment 2 involved algae alone as detailed in (Lamonica, Herbach, et al., 2016) (referred in that paper as “Experiment 1” in section 2.2.1.). Experiment 3 involved algae and duckweeds as detailed in (Lamonica, Herbach, et al., 2016) (referred in that paper as “Experiment 3” in section 2.2.2.).

### Experiments with cadmium

Experiment 4 involved algae and duckweeds, with two conditions in species composition: duckweeds alone, and algae and duckweeds together. We tested five different cadmium concentrations (0, 11.1, 20.2, 35.5 and 51.1 *μ*g/L) in triplicate for each condition. Three additional control beakers were inoculated with algae alone. The duration of this experiment was 14 days. From this experiment, we obtained different types of data under contaminant exposure: “monospecific data, duckweeds”, “two species data, duckweeds” and “two species data, algae”.

Experiment 5 involved the three species, with three conditions in species composition: duckweeds alone, algae and duckweeds, and algae, duckweeds and daphnids. We tested five different cadmium concentrations in triplicate for each condition (0, 2.25, 4.50, 6.88 and 9.09 *μ*g/L). The duration of this experiment was 21 days. From this experiment, we obtained the following data under contaminant exposure: “monospecific data, duckweeds”; “two species data, duckweeds” and “two species data, algae”; “complete microcosm data, duckweeds”, “complete microcosm data, algae” and “complete microcosm data, daphnids”.

Experiment 6 involved algae alone. We tested five different cadmium concentrations (0, 26.2, 36.4, 40.8 and 43.6 *μ*g/L) in triplicate. The duration of this experiment was 14 days. From this experiment, we obtained “monospecific data, algae”.

As mentioned in (Lamonica, Herbach, et al., 2016), we used measured cadmium concentrations in the medium instead of nominal ones. For that purpose, we measured dissolved cadmium concentrations as described by Clement et al. (BJP Clément et al., 2014) at days 2, 7, 14 (and day 21 for Experiment 5) in each beaker. We then calculated the arithmetic mean of all the measurements. In total, for Experiments 4, 5 and 6, we thus obtained 13 concentrations (0, 2.25, 4.50, 6.88, 9.09, 11.1, 20.2, 35.5, 51.1, 26.2, 36.4, 40.8 and 43.6 *μ*g/L) denoted by *C*_*j*_, *j* ∈ [0, 12] hereafter. The concentration in the controls of Experiments 4, 5 and 6 (that is with no contaminant) is denoted by *C*_0_, corresponding to index *j* = 0. This is also the case in Experiments 1 to 3, that were conducted without contaminant.

## Dynamic modelling

The description of the model follows the Overview, Design concepts and Details (ODD) protocol originally used for describing individual and agent-based models Grimm et al., 2010 but adapted here for a dynamic model based on Ordinary Differential Equations (ODE). The ODD protocol consists of seven elements. The first three elements provide an overview; the fourth element explains general concepts underlying the model’s design and the remaining three elements provide further details. Model code can be found at (Lamonica, Charles, et al., 2022a).

### Purpose

The model developed in this paper describes the dynamics of duckweeds, algae and daphnids under the microcosm conditions described in “Experiments and observed data” section. In particular, it aims at i) comparing the species dynamics both in isolation and together in order to highlight the interactions between the three species; and ii) describing the effects of cadmium on the different processes involved in the microcosm functioning. We first present the model of the three species’ dynamics without contaminant, then we show how we integrated cadmium effects in the model.

### Entities, state variables and scales

We model both duckweed and algal population dynamics but we only model two daphnid life history traits (growth and survival) that are involved in the interaction between algae and daphnids. The model involves five state variables. The two first ones refer to the numbers of algal cells per beaker in the two compartments of the microcosm at time *t* and cadmium concentration *C*_*j*_: the suspended algae in the water column (Compartment 1), denoted by *N*_1_(*t, C*_*j*_), and the settled algae at the bottom of the beaker (Compartment 2), denoted by *N*_2_(*t, C*_*j*_). The third state variable is the number of duckweed fronds per beaker at time *t* and cadmium concentration *C*_*j*_, denoted by *N*_*d*_(*t, C*_*j*_). The two other state variables refer to the daphnids: the number of alive daphnids in the microcosm through survival rate at time *t* and cadmium concentration *C*_*j*_, denoted by *S*(*t, C*_*j*_) and the daphnid size at time *t* and cadmium concentration *C*_*j*_, denoted by *L*(*t, C*_*j*_). The model is run on 21 days, corresponding to the duration of the longest experiment.

### Process overview and scheduling

Nine processes are modelled with a continuous time scale, using ODE. Two processes are related to intrinsic algal dynamics: settling of suspended algae and growth of both suspended and settled algae. One process is related to intrinsic duckweed dynamics: duckweed growth. One process concerns the algae-duckweed interaction with an interspecific competition. Two processes are related to daphnid life history traits: survival and growth. Two processes are related to algae-daphnid interaction: ingestion of algae by daphnids and location of daphnid for grazing. The last process is related to the effects of cadmium on the different parameters. An overall graphical representation of the implemented model is given in Figure S1 (Lamonica, Charles, et al., 2022c)

### Design concepts

#### Basic principles

The assumptions we make are based on the experimental design described in “Experiments and observed data” section. We assume that algae are uniformly distributed in the water column and at the bottom of the beaker at each time step and that the settling speed of suspended algae is constant throughout the water column. Therefore, the water volume occupied by the suspended algae is supposed to decrease at the same speed as algal settling. We assume that algae and duckweeds are competing only for nutrients in the medium.

We also assume that settled algae are too distant from duckweeds to interact with them, so that the interspecific competition only involves suspended algae. Interspecific competition has no effect on algae, as shown in Lamonica, Herbach, et al., 2016. We assume that cadmium affects the growth rates of all species, as well as competition intensity parameters and daphnid survival. Cadmium is supposed not to affect either the carrying capacities of algae and duckweeds or the algal settling rate.

#### Emergence

Algal and duckweed dynamics emerge both from their intrinsic dynamics (growth and settling for algae, growth for duckweeds) and from the interspecific competition between the two species. Algal dynamics also depends on daphnids through the quantity of algal cells that are consumed by daphnids. With cadmium, both dynamics emerge from the impact of cadmium on their respective growth and on the interaction.

#### Sensing

In order to determine the number of daphnids grazing in each compartment over time, we assume that daphnids, as pelagic species, preferentially feed in the water column Siehoff et al., 2009. We also assume that daphnids move to the sediment when the ratio of algal density in the water column over the bottom of the beaker is below a given threshold Siehoff et al., 2009.

#### Interactions

Intraspecific competition between algal cells and between duckweed colonies are taken into account in their respective logistic growth models. Algae and duckweeds interact through an interspecific competition process, described with a Lotka-Volterra type I interaction model. Algae and daphnids interact through a trophic relationship, namely grazing.

#### Stochasticity

We use stochasticity to describe variability on state variables, which sum up both uncertainties and variability sources within the processes. We suppose a normal distribution on the decimal logarithm of the number of algal cells per beaker in each compartment (in the water column and at the bottom of the beaker) Roger and Reynaud, 1978 and on the decimal logarithm of the number of duckweed fronds. For the number of daphnid survivors we consider a conditional binomial distribution Forfait-Dubuc et al., 2012 and a normal distribution for the daphnid size.

### Initialisation

As algae are inoculated in the water column only, the initial values for the number of algal cells per beaker in the water column and at the bottom of the beaker are 4 *×* 10^7^ and 0, respectively. The initial number of duckweed fronds is 8. The initial number of daphnids is 10, the initial survival rate is fixed to 1 (as all introduced daphnids are alive) and the initial daphnid size is drawn from a normal distribution (see hereafter section 4.1). As mentioned in section 2., we use measured cadmium concentrations 0, 2.25, 4.50, 6.88, 9.09, 11.1, 20.2, 35.5, 51.1, 26.2, 36.4, 40.8 and 43.6 *μ*g/L.

### Input data

The model does not use input data to represent time-varying environmental processes. Laboratory conditions are controlled and supposed to be constant over time.

### Submodels

All information on parameters and variables involved in the model are gathered together in SI Table S2 (Lamonica, Charles, et al., 2022c). Details about parameter estimation are given in “Statistical inference” section.

The deterministic part of algal dynamics in both compartments and of duckweed dynamics over time *t* (in days) is described with three coupled ODE. The deterministic part of daphnid survival and size are described with two other ODE that are presented in their integrated form.

#### Algae processes

We model the algae dynamics using logistic functions to describe algae growth in the water column and at the bottom of the beaker. We used an exponential decay of algal cells in the water column to describe sedimentation process.

#### Duckweed process

We model the duckweed growth using a logistic function.

### Daphnid processes

#### Survival

Survival rate at time *t* and cadmium concentration *C*_*j*_, *S*(*t, C*_*j*_), is described by an exponential decay with an instantaneous mortality rate, *m*_0_ (day^-1^), which is assumed to be time-independent Forfait-Dubuc et al., 2012:

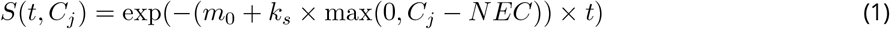

where *k*_*S*_ (*μ*g^-1^.L.day^-1^) represents the cadium effect intensity and *NEC* (No Effect Concentration) (*μ*g.L^-1^) is the concentration from which the contaminant has an effect on survival. When concentration *Cj* is below the *NEC*, max(0, *C*_*j*_ − *NEC*) is equal to 0, thus there is no effect on survival rate which only depends on natural mortality *m*_0_ and time *t*. However, when concentration *Cj* is superior to the NEC, max(0, *C*_*j*_ − *NEC*) is equal to the surplus of concentration and mortality due to cadmium is added to the natural mortality. We consider a conditional binomial stochastic model for *D*_*s*_(*t, C*_*j*_), the number of alive daphnids at time *t* and cadmium concentration *C*_*j*_ in the system Forfait-Dubuc et al., 2012:

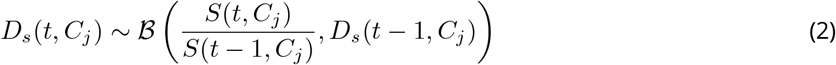

where *B* stands for the binomial law. For each concentration *C*_*j*_, the number of alive daphnids at time *t* depends on the number of alive daphnids at time *t* − 1 and on the survival probability between *t* − 1 and *t*, represented by 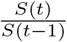.

We make here the implicit assumption that contaminant toxicokinetics is fast (which means that internal concentration in the organism is supposed to be equal to external concentration in the water *C*_*j*_) since cadmium have been shown to have a rapid toxicokinetic, especially a high capacity of bioaccumulation, at least in freshwater organisms (Gestin et al., 2021; Ratier and Charles, 2022).

#### Growth

Daphnid growth is described using a Von Bertalanffy growth model Von Bertalanffy, 1938. In addition, the daphnid size is supposed to follow a normal distribution with mean *L*(*t, C*_*j*_) and standard deviation *σ*_*L*_.

### Interaction processes

#### Interspecific competition process

We model the interspecific competition process between algae and duckweed using a unilateral Lotka-Volterra type I model, with an effect on duckweed dynamics only.

#### Ingestion process

The ingestion rate of a daphnid, *i*.*e*. the number of cells per beaker each daphnid consumes per day (denoted as *g*_1_(*t, C*_*j*_) in the water column and *g*_2_(*t, C*_*j*_) at the bottom of the beaker) is modelled with a Holling type II function of algal density in each compartment (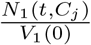 and 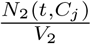), for a given daphnid

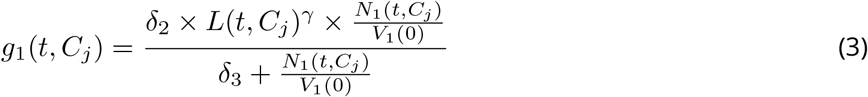

and

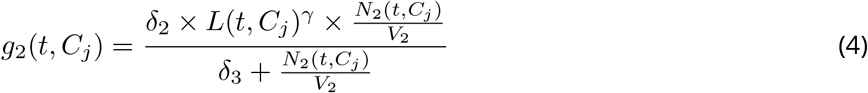

with δ_2_ (cell.daphnd^−1^.day^−1^.mm^−γ^) the maximum ingestion rate, *δ*_3_ (cell.mL^-1^) the algal density for which the ingestion rate is equal to half the maximum ingestion rate and *γ* (dimensionless) a regression coefficient.

#### Grazing location

The number of daphnids grazing in the water column at time *t* and cadmium concentration *C*_*j*_, *D*_1_(*t, C*_*j*_), is modelled with respect to the ratio *R*(*t, C*_*j*_) of algal density in compartment 1 over compartment 2 and the number of alive daphnids per beaker *D*_*s*_(*t, C*_*j*_):

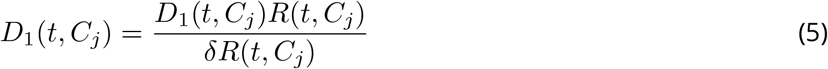

### Cadmium effects

We suppose that the survival process is affected by cadmium according to Eq.(5).

We suppose that only growth rates and parameters of competition intensity are affected by cadmium, as already assumed in Lamonica, Herbach, et al., 2016. We choose a three-parameter log-logistic function to describe the effect of cadmium at concentration *C*_*j*_ on each affected parameter *p*:

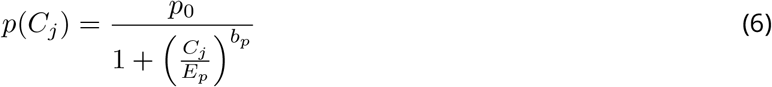

where *p*_0_ is the value of parameter *p* in the control, *E*_*p*_ is the cadmium concentration at which 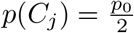, which is equivalent to an *EC*_50_, and *b*_*p*_ is the curvature coefficient of the log-logistic function.

### Complete model

Finally, the deterministic part of the model describing the functioning of the whole microcosm is expressed as follows:

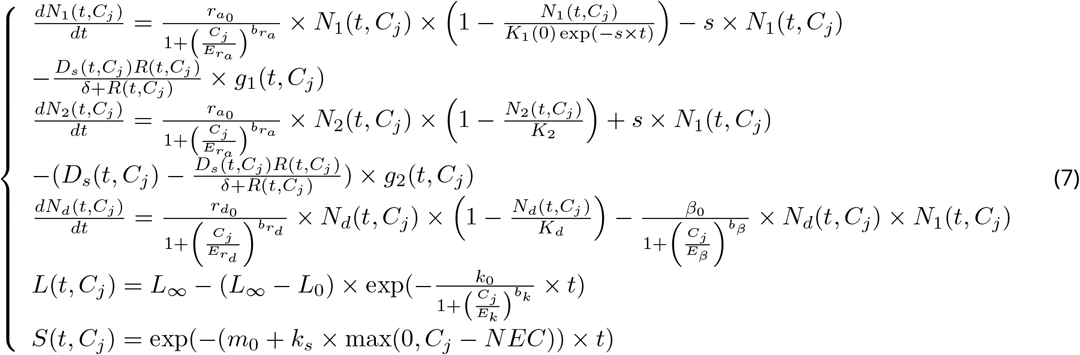

The same model can be applied when daphnids are absent, by setting *D*_*s*_(*t, C*_*j*_) = 0 and thus *D*_1_(*t, C*_*j*_) = 0. In this case, the two last equations must also be removed. The same model can be applied when duckweeds are absent, by setting *N*_*d*_(*t, C*_*j*_) = 0 and removing the third equation. The same model can be applied when algae are absent, by setting *N*_1_(*t, C*_*j*_) = 0 and removing the first two equations. When the water column is stirred (*i*.*e*. when algae are supposed not to settle and duckweeds and daphnids are absent), the settling rate *s* is assumed to be zero and the second and two last equations must be removed.

At each time step, the decimal logarithm of the number of algal cells per beaker in the water column follows a normal distribution of mean *N*_1_(*t, C*_*j*_) and standard deviation 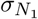. The decimal logarithm of the number of algal cells per beaker on the sediment follows a normal distribution of mean *N*_2_(*t, C*_*j*_) and standard deviation 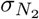. The decimal logarithm of the number of duckweed fronds per beaker follows a normal distribution of mean *N*_*d*_(*t, C*_*j*_) and standard deviation 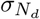.

## Statistical inference

In order to check if our model satisfactorily described the microcosm functioning, we used Bayesian inference to fit the model simultaneously to all our experimental data from the six above mentioned experiments. Estimates obtained for all the parameters are called “reference estimates” hereafter.

### Parameter prior distributions

We defined prior distributions summarising all information on each parameter available in advance (SI Table S2 (Lamonica, Charles, et al., 2022c)). Some of the prior distributions described the decimal logarithm of the parameter because of an expected large range of possible values (for instance, 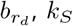 and *β*_0_) or extreme orders of magnitude (*e*.*g*., large or small). Other prior distributions were defined based on previous experiments that were conducted using the same experimental device (Billoir, Delhaye, B Clément, et al., 2011; Billoir, Delhaye, Forfait, et al., 2012; Delhaye, 2011; Lamonica, B Clément, et al., 2016) 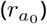, or based on additional experiments specifically conducted in the laboratory 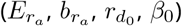. At last, the remaining distributions were based on literature values (Billoir, Delignette-muller, et al., 2008; Biron et al., 2012; DeMott, 1982; Egloff and Palmer, 1971) (*k*_0_), except for parameters on which we had very vague information 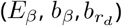, so that their prior distributions were chosen as flat.

### Computation

Monte Carlo Markov Chain (MCMC) computations were performed using the JAGS software via the *rjags* R package (Plummer, 2009; Team, 2013), after the model was discretised using the Euler method with a time step equal to 0.1 as stated in (Lamonica, B Clément, et al., 2016). Three chains were run. A total of 20000 iterations was performed as a burn-in phase and inference was based on 100000 additional iterations for each of the three chains. To check the convergence of the estimation process, we used the Gelman and Rubin convergence diagnostic (Gelman and Rubin, 1992) with a cut-off of 1.01. R code can be found at (Lamonica, Charles, et al., 2022a).

### Posterior Predictive Check

To check posterior predictions of the model, we simulated new date at all experienced time steps and tested concentrations taking into account parameter uncertainties and stochasticity of the model (Lamonica, B Clément, et al., 2016). 95% of the observed data are expected to be contained in the 95% credibility band of the predicted data, got from 2.5% and 97.5% percentiles of the predictions. R code can be found at (Lamonica, Charles, et al., 2022a).

### Look-back on the experimental design

We aim at determining which types of data could be sufficient to accurately (in terms of mode of the posterior distribution) and precisely (in terms of dispersion of the posterior distribution) estimate parameters of stress functions for the different species (table 1). Our reference was the posterior distributions obtained when estimating parameters from the whole dataset, considering them as the “best possible estimates in the present case study in view of the model and all available data”. To evaluate the information provided by certain data, we built partial datasets by removing these data from the whole dataset. Then, we re-estimated the model parameters using these partial datasets and compared the newly obtained estimates with the reference ones.

**Table 1.** Parameters of interest.

| Symbol | Definition | Unit | Prior distribution*<br>or value | Sources** | 2.5, 50, 97.5 %<br>posterior quantiles |
| --- | --- | --- | --- | --- | --- |
| $r_{a0}$ | Intrinsic algal growth rate in the control | day<sup>-1</sup> | $\mathcal{U}(0,2)$ | [1] | 0.78 0.83 0.87 |
| $E_{ra}$ | The concentration at which $r_{a0}$ is reduced by 50 % | $\mu\text{g.L}^{-1}$ | $\log_{10}(E_{ra}) \sim \mathcal{N}(1.78, 0.1)$ | [2] | 1.54 1.56 1.58 |
| $b_{ra}$ | Curvature coefficient | - | $\log_{10}(b_{ra}) \sim \mathcal{N}(0.24, 0.1)$ | [2] | 0.37 0.43 0.49 |
| $\beta_0$ | Competition intensity of algae on duckweeds in the control | # of cells per beaker<sup>-1.day<sup>-1</sup></sup> | $\log_{10}(\beta_0) \sim \mathcal{U}(-11, -9)$ | [2] | -9.53 -9.46 -9.40 |
| $E_\beta$ | The concentration at which $\beta_0$ is reduced by 50 % | $\mu\text{g.L}^{-1}$ | $\log_{10}(E_\beta) \sim \mathcal{U}(-1, 4)$ | Vague | -0.99 -0.77 -0.15 |
| $b_\beta$ | Curvature coefficient | - | $\log_{10}(b_\beta) \sim \mathcal{U}(-3, 3)$ | Vague | -1.11 -0.82 -0.62 |
| $m_0$ | Daphnid mortality rate in the control | daphnid.day<sup>-1</sup> | $\log_{10}(m_0) \sim \mathcal{U}(-4, 0)$ | Vague | -1.89 -1.71 -1.56 |
| $k_S$ | Slope of survival stress function | $\mu\text{g}^{-1}.\text{L.day}^{-1}$ | $\log_{10}(k_S) \sim \mathcal{U}(-4, 4)$ | Vague | -1.80 -1.54 -1.30 |
| $NEC$ | No Effect Concentration on daphnid survival | $\mu\text{g.L}^{-1}$ | $\log_{10}(NEC) \sim \mathcal{U}(-2, 1)$ | Vague | 0.47 0.65 0.76 |
| $k_0$ | Daphnid growth rate in the control | day<sup>-1</sup> | $\mathcal{N}(0.11, 0.030)$ | [3] | 0.138 0.146 0.154 |

| Symbol | Definition | Unit | Prior distribution*<br>or value | Sources** | 2.5 % | 50 % | 97.5 %<br>posterior quantiles |
| --- | --- | --- | --- | --- | --- | --- | --- |
| $E_k$ | The concentration at which $k_0$ is reduced by 50 % | $\mu\text{g.L}^{-1}$ | $\log_{10}(E_k) \sim \mathcal{U}(-1,4)$ | | 1.26 | 1.68 | 3.31 |
| $b_k$ | Curvature coefficient | - | $\log_{10}(b_k) \sim \mathcal{U}(-3,3)$ | | -0.68 | -0.25 | 0.015 |
| $r_{d0}$ | Intrinsic duckweed growth rate in the control | $\text{day}^{-1}$ | $\mathcal{U}(0,2)$ | [2] | 0.23 | 0.24 | 0.25 |
| $E_{r_d}$ | The concentration at which $r_{d0}$ is reduced by 50 % | $\mu\text{g.L}^{-1}$ | $\log_{10}(E_{r_d}) \sim \mathcal{N}(2.44, 0.2)$ | [2] | 1.96 | 2.30 | 2.67 |
| $b_{r_d}$ | Curvature coefficient | - | $\log_{10}(b_{r_d}) \sim \mathcal{U}(-3,3)$ | Vague | -1.05 | -0.91 | -0.77 |

\*Prior distribution: $\mathcal{N}$ stands for the normal law, $\mathcal{U}$ stands for the uniform law. \*\*Sources: [1] (Lamonica, B Clément, et al., 2016), [2] (Lamonica, Herbach, et al., 2016), [3] (Billoir, Delignette-muller, et al., 2008).

Four partial datasets (numbered A to D) were used to estimate the stress function parameters. They are summarized in table 2. We evaluated only information provided by data collected under contaminated conditions, removing them successively to build the partial datasets. Regarding data without contaminant, including the controls in Experiments 4 to 6, they were kept in all partial datasets.

**Table 2.** Experimental design and partial datasets. The black dashed, gray dashed, black and gray rectangles refer to the data that have been removed from the entire dataset to build partial dataset A, B, C and D, respectively.

| Cadmium concentrations | Monospecific data | Two species data | Complete microcosm data |
| --- | --- | --- | --- |
| $C_0$ | Algae<br>Duckweeds | Algae, duckweeds<br>Algae, daphnids | Algae, duckweeds, daphnids |
| $C_{1-4}$ | Algae<br>Duckweeds | Algae, duckweeds | Algae, duckweeds, daphnids |
| $C_{5-8}$ | Algae<br>Duckweeds | Algae, duckweeds | |
| $C_{9-12}$ | Algae | | |

Dataset A included all data except the “monospecific data” with contaminant. This corresponds to exclude data collected from beakers containing only one species. Thus, data from Experiment 6 and data “duckweeds alone” from Experiments 4 and 5 were not included in dataset A. Dataset B included all data except the “two species data” with contaminant. This corresponds to exclude data collected from beakers with both duckweeds and algae. Thus, duckweed and algae data collected from beakers containing both duckweeds and algae from Experiments 4 and 5 were not included in dataset B. In dataset C, we only included the “complete microcosm data”, which corresponds to include only data collected from Experiment 5 with the three species. Dataset D was used to evaluate the information provided by data collected at concentrations lower than 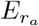 and 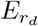 (*EC*_50_ values) for algae and duckweeds, respectively. Thus, dataset D included all data except those related to duckweeds and algae exposed to the lowest cadmium concentrations (“monospecific data” and “two species data”) collected from Experiment 5. All in all, we tested the minimum necessary dataset in terms of species combinations (one, two, or three species) from datasets A, B and C, and of concentration range via dataset D.

## Results

### Model fit and parameter estimates

Our MCMC algorithm always consistently converged according to Gelman and Rubin diagnostics for each simulation. The corresponding 2.5%, 50% and 97.5% quantiles of the posteriors for parameters of interest are summarised in table 1. To keep results clear enough, we only display fitting results from data of Experiment 5 (measured cadmium concentrations of 0, 2.25, 4.50, 6.88 and 9.09 *μ*g/L) as medians of the credibility band for predicted data on algae dynamics (figure 2) and daphnid survival (figure 3). On a general point of view, data were satisfactorily described by the model, with between 91% and 98% of observed data encompassed in the 95% credibility band of the predictions for the different species.

**Figure 2.**
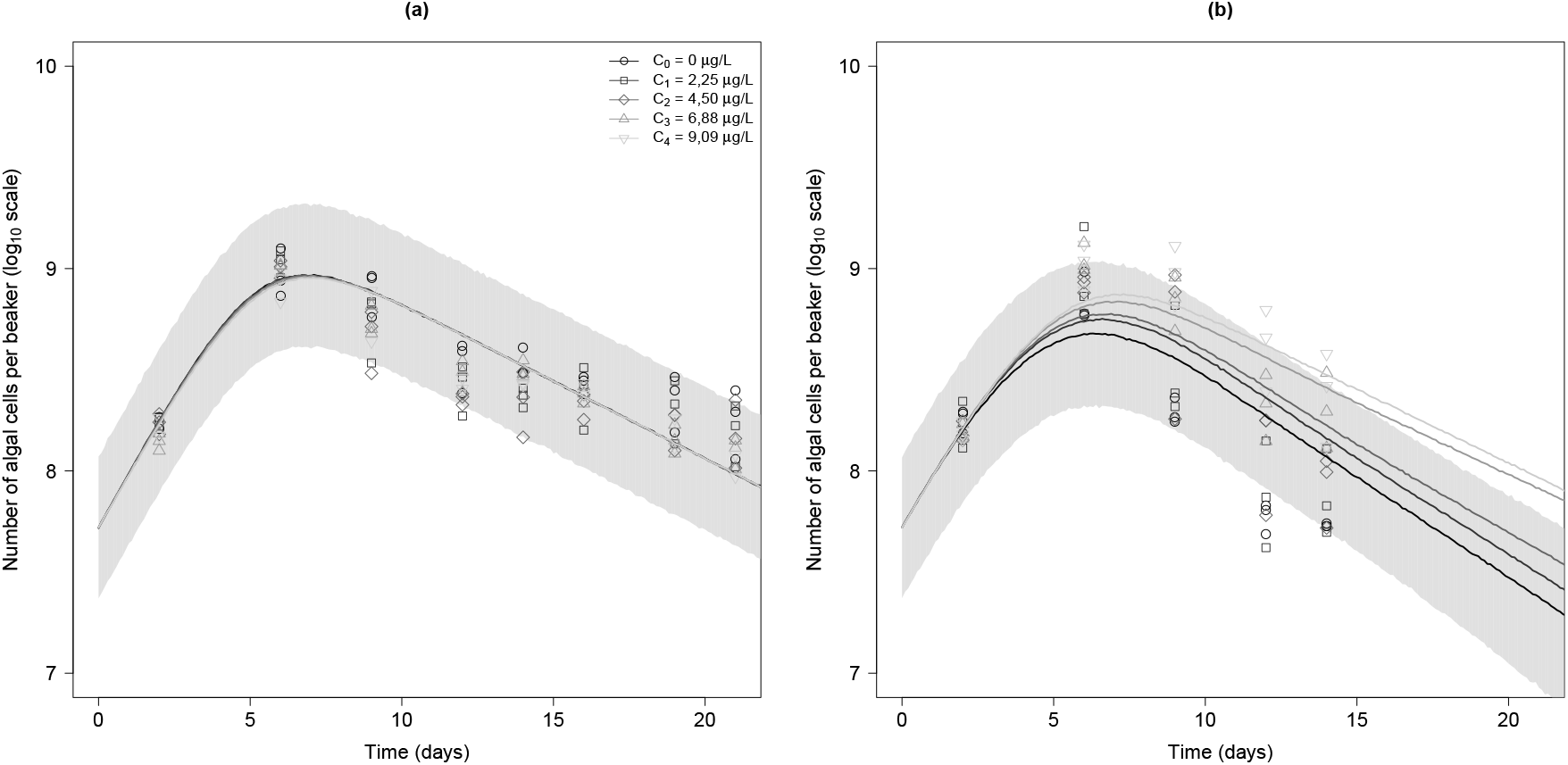
Model fitting for state variables related to algae dynamics: (a) with duckweeds; (b) with duckweeds and daphnids. Symbols refer to the different tested cadmium concentrations of Experiment 5. Plain lines stand for the fitted model with each parameter equal to its median value at each concentration *C*_*j*_. The light grey area corresponds to the 95% credible band of the predicted data in the control.

**Figure 3.**
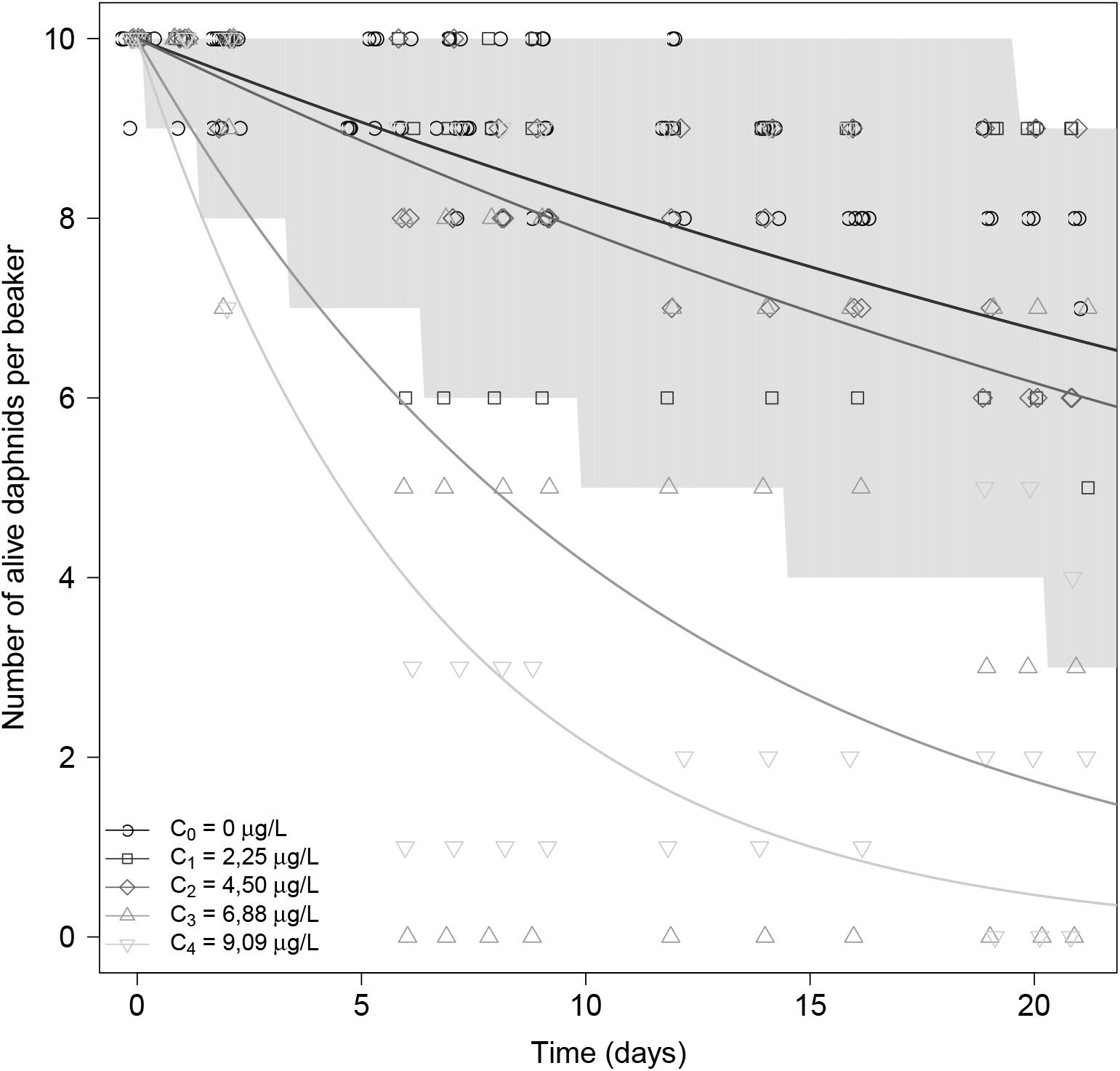
Fit plot for the number of alive daphnids per beaker. Symbols refer to the different tested cadmium concentrations of Experiment 5. Plain lines stand for the fitted model with each parameter equal to its median value at each concentration *C*_*j*_. The light grey area corresponds to the 95% credible band of the predicted data in the control.

Marginal posterior distributions of the estimated parameters are shown in SI (Figure S4 (Lamonica, Charles, et al., 2022c)). We obtained narrow posterior distributions for almost all parameters, in particular parameters of interest, with the exception of parameters related to algae-daphnids interaction (grazing). The narrowness of posterior distributions indicates that sufficient information was available in the data to get posterior distributions of model parameters that are more precise than their priors. Such a gain of knowledge makes us confident in our fitting process.

#### Algae dynamics and daphnid survival

In the presence of duckweeds only (figure 2(a), control), the number of algal cells per beaker in the water column increased during the first seven days when growth is higher than settling, and then decreased as growth declined while settling was continuing. In the presence of daphnids plus duckweeds, the global algal dynamics in the control was similar to the one without daphnids; however the number of algal cells per beaker was lower, due to daphnid grazing (figure 2(b), control). There was additionally no effect of cadmium on the algal dynamics when duckweeds were present (figure 2(a)). However, with daphnids, differences between tested concentrations appeared from the sixth day of experiment: the higher the cadmium concentration, the higher the number of algal cells (figure 2(b)). This may be due to the decrease in daphnid number (figure 3), daphnid survival being highly affected by cadmium, particularly at the two highest tested concentrations *C*_3_ and *C*_4_.

#### Look-back on the experimental design

To take into account the potential effect of correlations between parameters, we compared 95% credibility intervals of the stress functions predicted from the joint posterior distributions obtained for each partial dataset (A to D) to the one obtained from the whole dataset (figure 4). The 95% credibility intervals of the predicted stress functions on daphnid growth rate and survival for each partial dataset appear superimposed to the 95% credibility intervals of the predicted stress functions for all the data. This result was expected since daphnid’s survival and size data were included in all datasets, the whole and partial ones.

**Figure 4.**
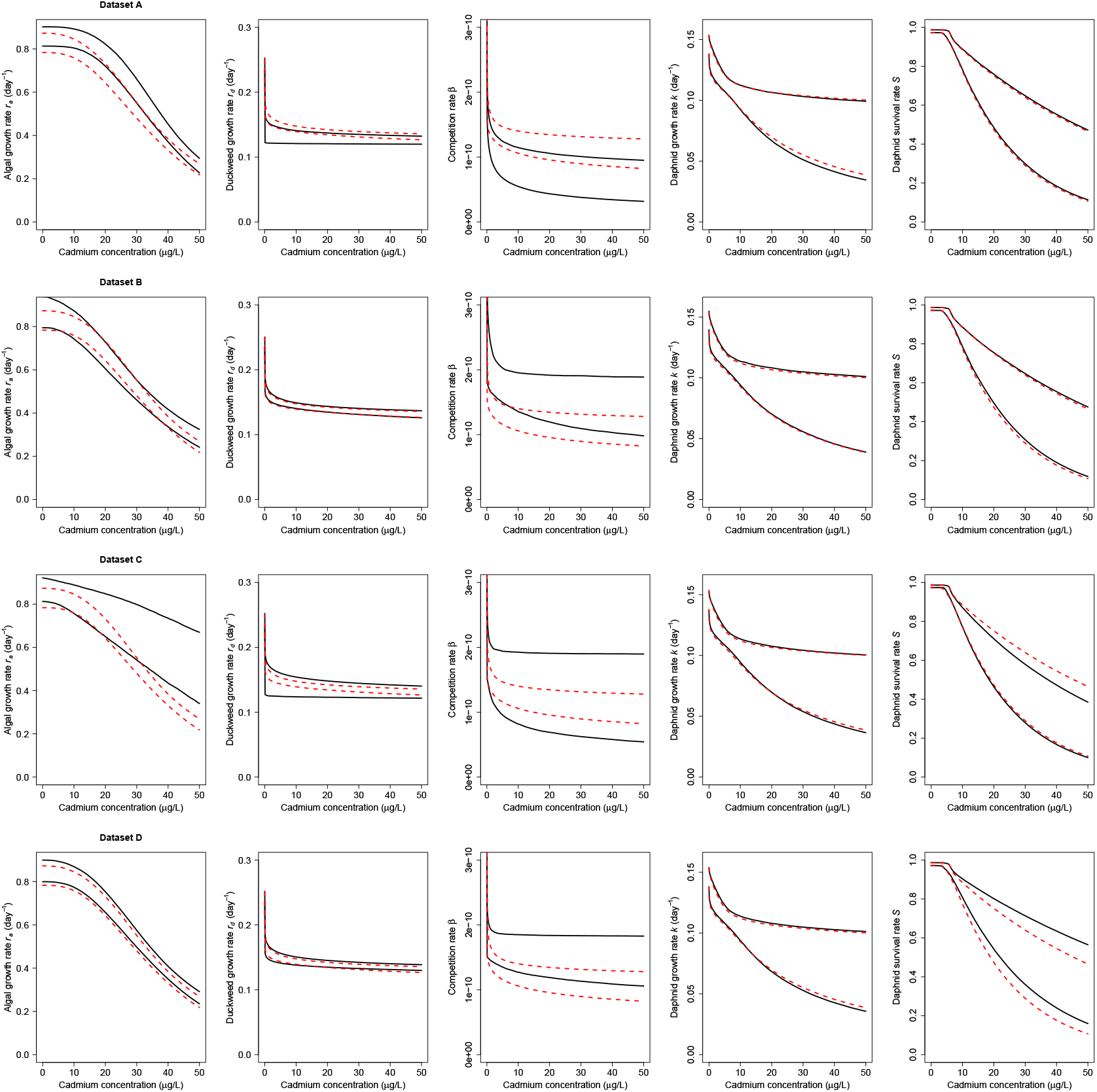
Comparison of stress functions obtained with complete and partial datasets. Plain black lines delimit the 95% credible bands for the stress functions obtained with the partial dataset, while red dotted lines delimit the 95% credible bands for the stress functions obtained with the complete dataset.

When “monospecific” data were removed (dataset A), the predicted stress functions were different from the reference ones with larger 95% credibility bands, particularly for competition parameter *β*. When “two species” data were removed (dataset B) the predicted stress functions for both algal and duckweed growth rates (*r*_*a*_ and *r*_*d*_) were very close to the reference ones. On the contrary, the predicted stress function for competition parameter *β* was overestimated, and showed more uncertainty. When “monospecific” and “two species” data were removed (dataset C), we obtained very large 95% credibility intervals for predicted stress functions for both growth rates and the competition parameter. At last, dataset D (without data related to duckweeds and algae exposed to the lowest concentrations *C*_1_ to *C*_4_) led to very similar predicted stress functions for both algal and duckweed growth rates (*r*_*a*_ and *r*_*d*_) compared to the reference ones, while for competition parameter *β* the predicted stress function was overestimated with a greater uncertainty.

## Discussion

### Cadmium effect

#### On parameters and processes

For parameters related to cadmium effect on algae and duckweeds, we obtained similar estimates to the ones obtained in a previous study involving only these two species (Lamonica, Herbach, et al., 2016). However, posterior distributions were narrower in the present study for some of the parameters, e.g. 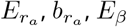 and *b*_*β*_, mainly thanks to additional data we considered for fitting.

Parameters of the stress function on daphnid survival showed narrow posterior distributions. Parameter *NEC* was estimated at 4.47 [2.95, 5.75] *μ*g/L while other authors estimated either higher *NEC* values (8.6 *μ*g/L (Nebeker et al., 1986)) or lower ones (0.720 [0.0427, 1.78] (Forfait-Dubuc et al., 2012)) from monospecific studies. Our *NEC* estimate was high compared to the one obtained with data from the same microcosm, but including five species and sediment, namely 1.8 [1.2, 2.3] *μ*g/L (Billoir, Delhaye, Forfait, et al., 2012). Nevertheless, the *NEC* estimated by Billoir et al. was only based on survival data, and thus did not include data related to the dynamics of the other species. In particular, the algae dynamics link to the number of surviving daphnids was ignored. We thus also estimated the *NEC* only using survival data to compare 95% credible interval to the one of (Billoir, Delhaye, Forfait, et al., 2012). We obtained a *NEC* value of 3.47 [0.030, 5.50] *μ*g/L. This credible interval is quite large because the number of alive daphnids per beaker was highly variable, but it contains the credible interval obtained by (Billoir, Delhaye, Forfait, et al., 2012). The number of alive daphnids per beaker revealed difficult to describe because of the high inter-replicate variability between the tested cadmium concentrations.

In the literature, the effects of cadmium on daphnid growth may vary a lot from one study to another: *EC*_10_ = 7.3 *μ*g/L for 17 days in(Knops et al., 2001) (monospecific bioassay conditions), *NEC* = 0.15 *μ*g/L in (Billoir, Delhaye, Forfait, et al., 2012) (five-species microcosm conditions) or *EC*_50_ = 2.7 *μ*g/L for 21 days in (BJP Clément et al., 2014) (five-species microcosm conditions). In the present study, such effects are expressed through both parameters *b*_*k*_ and *E*_*k*_ (*i*.*e*., *EC*_50_). We obtained a very high but imprecise estimate for *E*_*k*_ (47.9 [18.2, 2042] *μ*g/L) indicating that daphnids were less sensitive in our experiment than in the one conducted by (BJP Clément et al., 2014). However, the low values of curvature coefficient *b*_*k*_ (0.56 [0.21, 1.04] *μ*g/L) indicated that daphnid growth rate was already affected at the lowest concentrations, as also mentioned by (Billoir, Delhaye, Forfait, et al., 2012).

#### On the functioning of the microcosm

The microcosm functioning with cadmium or not is summarised in figure 5. Cadmium effects on daphnid processes corresponded to a negative direct effect on survival, in particular at concentrations *C*_3_ and *C*_4_ (figure 3), as well as to a lighter negative direct effect on growth (SI, Figure S3 (Lamonica, Charles, et al., 2022c)). These direct effects of cadmium on daphnid processes impact both algae and duckweeds. Indeed they induced a decrease in daphnid grazing that led to an increase in algal density with increasing cadmium concentrations. Such a result was supported by the absence of a cadmium effect on algal growth at concentrations below 10 *μ*g/L, that was a positive indirect effect of cadmium on algae below 10 *μ*g/L. In addition, there was a negative direct effect of cadmium on the competition intensity. This latter did not compensate the negative direct effect of cadmium on growth of duckweeds, especially since algal density became higher, leading to a decrease in duckweed density.

**Figure 5.**
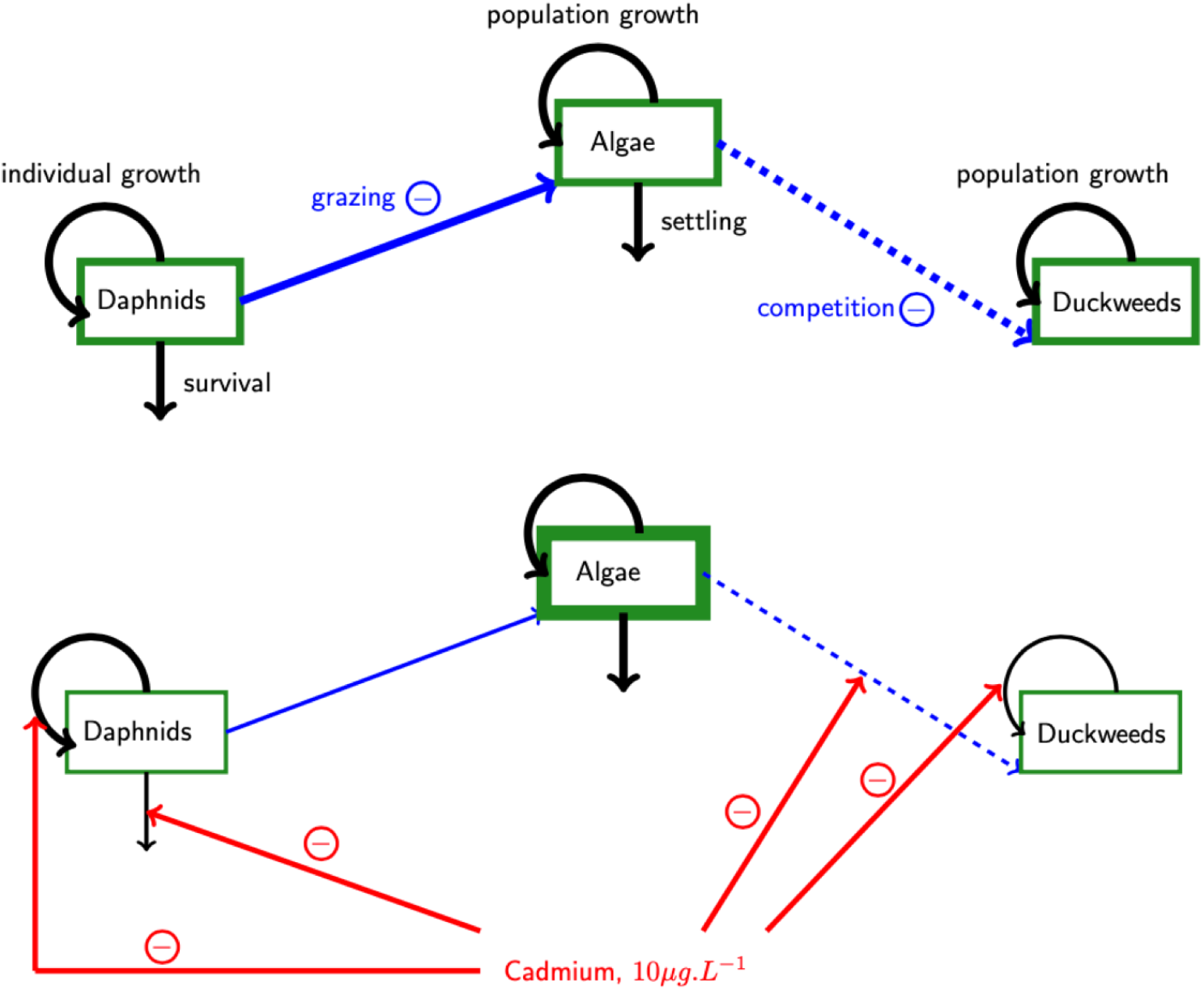
Global perturbation of the microcosm functioning without cadmium (above) and with cadmium (10 *μ*g/L, below). Green boxes represent species population, black arrows represent species processes, black arrows represent interactions between species, red ligthnings represent cadmium effect on the different processes. Thickness of boxes lines is proportional to population size and thickness of arrows is proportional to process intensity.

Deciphering the cascade of cadmium effects on the three species is finally possible thanks to our modelling approach coupled with experiments. Unravelling the chemical direct and indirect effects as well as the interactions between species is necessary to correctly interpret the global effect of a chemical substance on the functioning of a species community. Nevertheless, it is much more challenging when the number of species is increasing (Lamonica, Herbach, et al., 2016). Hence, the model fits of the present study were overall satisfactory although some predicted data were overestimated compared to observed ones. In particular, the number of algal cells per beaker in the presence of both duckweeds and daphnids were overestimated by our model, as well as the number of duckweed fronds per beaker with algae and daphnids (SI, Figure S2 (b) (Lamonica, Charles, et al., 2022c)).

#### Look-back on the experimental design

In ecotoxicology, optimising the experimental designs is not a recent concern (Albert et al., 0001; Andersen et al., 2000; Forfait-Dubuc et al., 2012; Wright and Bailer, 2006), but today mainly relates to the increasing use of concentration-response or effect models (Chèvre and Brazzale, 2008; Forfait-Dubuc et al., 2012; Holland-Letz and Kopp-Schneider, 2015; Keddig et al., 2015; Khinkis et al., 2003; Sitter and Torsney, 1995; Wang et al., 2006). As formal optimisation first applied to monospecific bioassays, it mainly focused on the tested concentrations (range and number of concentrations) and on the number of tested individuals (per tested concentration and in total) (Forfait-Dubuc et al., 2012). Formal optimisation is less suitable for microcosm experiments because microcosms are more complex devices. Moreover, microcosms are not standardised experimental tools since they are usually set up on a case-by-case basis, according to the specific objectives of the study (Cairns Jr and Cherry, 1997; Crossland and La Point, 1992). Thanks to our modelling approach, we were able to question the relevance of using certain data to estimate the chosen parameters. Hence, using dataset B provided stress functions similar to the reference ones based on all available data, even though there were changes in individual parameters, which needs to be taken into account when aiming at estimating *EC*_50_ in particular. We also showed that stress function on the interspecific competition parameter (*β*) from partial dataset D differed from the reference one, while the stress functions on the processes related to growth (for both algae and duckweeds) remained unchanged. Such results suggest that two-species data are not fully necessary, while a larger range of tested concentrations would be strongly recommended to estimate parameters related to effects on interactions between species.

Datasets A and B were chosen in order to test if monospecific, respectively two-species, microcosms were necessary to estimate parameters, and dataset C was chosen to test if the complete microscosm alone was sufficient to estimate parameters. Similarly, dataset D was chosen to test if reducing the number of tested concentrations would affect the quality of parameter estimates. Omitting the different species combinations, as well as reducing the number of tested concentrations, would save a lot of time and experimental effort, especially since adding one concentration to the design implies adding the number of replicates times the number of species combination (and not only the number of replicates). Also, cutting off some of the species combinations or some of the tested concentrations would permit to increase the number of replicates per treatment. More replicates may help capture the variability of the system, allowing to better take uncertainties into account. In addition, when using animal species, the overall objective is to reduce the number of organisms involved in experiments.

Even if it remains difficult to know *a priori* which types of data would be absolutely necessary to best estimate stress function parameters, the pragmatic look-back we performed using modelling may guide further experiments with microcosms, for instance for other contaminants effects, dynamics under modified abiotic conditions, or even other species interactions. In particular, our study suggests that experiments can be specifically selected to gain knowledge on a three-species microcosm. In the end, we could make the following recommendations for further ecotoxicological studies with a microcosm device: a first experiment with the complete microcosm only (*i*.*e*., with the three species) and a tested concentration range limited by the sensitivity of the most sensitive species; then a second experiment with monospecific microcosms only and a tested concentration range limited by the sensitivities of the two less sensitive species. If needed, additional experiments without contaminant may involve different combinations of species depending on their connections to each others. More generally, we would suggest that collecting data of monospecific and complete microcosms with contaminant might be sufficient to assess the contaminant effects, as long as in-depth knowledge of the functioning without contaminant is available. Nevertheless, there are limits to how transferable those recommendations are. When using another microcosm with different species or additional species, interactions between species still need to be properly investigated without contaminant first. Some of the species combinations with contaminant may not be discarded according to the direction or type of the interactions between those species. When using different contaminant, especially contaminants with different mode of action, discarding the lowest concentrations might be an issue. For instance, endocrine disrupting contaminants may show a strong non-linear effect, including effects at low concentrations; in that case scenario low concentrations should obviously be maintained in the experimental design.

## Conclusion and perspectives

We provided *EC*_50_ values for the different processes affected by cadmium. Thanks to the understanding of the underlying processes that occurred in the microcosm functioning, we also managed to identify the cascade of cadmium effects induced by the interactions between species. In addition, we got back from modelling to experiments in order to determine which of the collected data were necessary and sufficient to precisely estimate model parameters, leading us to suggest a more efficient experimental design. Finally, we (1) highlighted the importance of interactions by identifying the effect cascade occurring within a small ecosystem under chemical pressure; and (2) showed that alternative use of experimental data can help conceiving experimental designs for a microcosm study.

Our method also permitted to assess which data to include when estimating parameters of interest in a dynamic ecosystem model from a laboratory based microcosm ecotoxicity study. Such an approach could be enhanced to better foresee further experiments with microcosms based a similar model. Beyond this, if parameters are simultaneously estimated a whole dataset, this makes possible to compare these reference estimates with those obtained with partial datasets. This gives knowledge on the data dependency in the modelling results. Last but not least, such a retrospective and descriptive sensitivity analysis puts light in the fact that data quality and design are more beneficial for modelling purpose than quantity. Ideally, as the use of models and big data in ecology increases (Van Den Brink et al., 2016), modellers and experimenters could collaboratively and profitably elaborate model-guided experiments.

## Acknowledgements

Authors thank Pauline Le Quellec et Ludovik Hauduroy for their contribution to the experiments. Preprint version 6 of this article has been peer-reviewed and recommended by Peer Community In Ecotoxicology and Environmental Chemistry (https://doi.org/10.24072/pci.ecotoxenvchem.100002).

## Fundings

Financial support from ENTPE (École Nationale des Travaux Publics de l’État) and IXXI institute (Institut Rhô-nalpin des Systèmes Complexes).

## Conflict of interest disclosure

The authors declare that they comply with the PCI rule of having no financial conflicts of interest in relation to the content of the article.

## Data, script, code, and supplementary information availability

Data are available online: https://doi.org/10.5281/zenodo.6598408 Script and codes are available online: https://doi.org/10.5281/zenodo.6598408 Supplementary information is available online: https://doi.org/10.5281/zenodo.6598408

## Notes

https://doi.org/10.5281/zenodo.6598408

### Competing Interest Statement

The authors have declared no competing interest.

### Summary of Updates

This version of the article has been peer-reviewed and recommended by Peer Community In Ecotoxicology and Environmental Chemistry.

