## supplementary information (figures + tables) for "Chemical effects on ecological interactions within a model-experiment loop" (from another version of this preprint: version 5, 2022-09-13)

### Supporting information

#### 1 Tables

Table 2: Table S2: Variables and parameters of the model.

| Symbol | Definition | Unit | Prior distribution* or value | Sources** | 2.5, 50, 97.5 % posterior quantile[illegible] | | |
|---|---|---|---|---|---|---|---|
| $N_1(t, C_j)$ | Number of algal cells per beaker in the water column at time $t$ and concentration $C_j$ | # of cells per beaker | | | | | |
| $N_2(t, C_j)$ | Number of algal cells per beaker at the bottom at time $t$ and concentration $C_j$ | # of cells per beaker | | | | | |
| $V_1(0)$ | Volume of the water column at $t = 0$ | mL | 2000 | | | | |
| $r_{a_0}$ | Intrinsic algal growth rate in the control | day$^{-1}$ | $\mathcal{U}$ (0,2) | [1] | 0.78 | 0.83 | 0.8[illegible] |
| $E_{r_a}$ | The concentration at which $r_{a_0}$ is reduced by 50 % | $\mu$g.L$^{-1}$ | $\log_{10}(E_{r_a}) \sim \mathcal{N}$ (1.78,0.1) | [2] | 1.54 | 1.56 | 1.5[illegible] |
| $b_{r_a}$ | Curvature coefficient | - | $\log_{10}(b_{r_a}) \sim \mathcal{N}$ (0.24,0.1) | [2] | 0.37 | 0.43 | 0.4[illegible] |
| $K_1(0)$ | Carrying capacity in the water column for algae at $t = 0$ | # of cells per beaker | $\log_{10}(K_1(0)) \sim \mathcal{U}$ (8,11) | [1] | 9.54 | 9.59 | 9.6[illegible] |
| $K_2$ | Carrying capacity for algae at the bottom | # of cells per beaker | $\log_{10}(K_2) \sim \mathcal{U}$ (8,11) | [1] | 9.12 | 9.22 | 9.3[illegible] |
| $s$ | Settling rate | day$^{-1}$ | $\log_{10}(s) \sim \mathcal{U}$ (-2,0) | [1] | -0.78 | -0.75 | -0.[illegible] |

| Symbol | Definition | Unit | Prior distribution* or value | Sources** | 2.5, 50, 97.5 % posterior quantiles | | |
|---|---|---|---|---|---|---|---|
| $\beta_0$ | Competition intensity of algae on duckweeds in the control | # of cells per beaker$^{-1}$.day$^{-1}$ | $\log_{10}(\beta_0) \sim \mathcal{U}$ (-11,-9) | [2] | -9.53 | -9.46 | -9.[illegible] |
| $E_\beta$ | The concentration at which $\beta_0$ is reduced by 50 % | $\mu$g.L$^{-1}$ | $\log_{10}(E_\beta) \sim \mathcal{U}$ (-1,4) | Vague | -0.99 | -0.77 | -0.[illegible] |
| $b_\beta$ | Curvature coefficient | - | $\log_{10}(b_\beta) \sim \mathcal{U}$ (-3,3) | Vague | -1.11 | -0.82 | -0.[illegible] |
| $g_1(t, C_j)$ | Number of cells per beaker each daphnid consumes per day in the water column at time $t$ and concentration $C_j$ | # cell.day$^{-1}$.daphnid$^{-1}$ | | | | | |
| $g_2(t, C_j)$ | Number of cells per beaker each daphnid consumes per day at the bottom of the beaker at time $t$ and concentration $C_j$ | # cell.day$^{-1}$.daphnid$^{-1}$ | | | | | |
| $\gamma$ | Regression coefficient | - | 3 | [1] | | | |
| $\delta_2$ | Maximum ingestion rate | cell.daphnid$^{-1}$.day$^{-1}$.mm$^{-3}$ | $\log_{10}(\delta_2) \sim \mathcal{U}$ (5,7.1) | [3,4,5,6] | 6.55 | 6.99 | 7.[illegible] |
| $\delta_3$ | Algal density at which maximum ingestion rate is reduced by 50% | cell.mL$^{-1}$ | $\log_{10}(\delta_3) \sim \mathcal{U}$ (5.5,7) | [3,4,5,6] | 5.86 | 6.36 | 6.[illegible] |
| $D_1(t, C_j)$ | Number of alive daphnid in the water column per beaker at time $t$ and concentration $C_j$ | # daphnids | | | | | |
| $\delta$ | Ratio of algal densities for which half of living daphnids graze in the water column | - | $\log_{10}(\delta) \sim \mathcal{U}$ (-3,3) | Vague | -3.14 | -2.83 | -2.[illegible] |

| Symbol | Definition | Unit | Prior distribution* or value | Sources** | 2.5, 50, 97.5 % posterior quantiles | | |
|---|---|---|---|---|---|---|---|
| $S(t, C_j)$ | Daphnid survival probability at time $t$ and concentration $C_j$ | - | | | | | |
| $D_s(t, C_j)$ | Number of alive daphnid per beaker at time $t$ and concentration $C_j$ | # daphnids | | | | | |
| $m_0$ | Daphnid mortality rate in the control | daphnid.day$^{-1}$ | $\log_{10}(m_0) \sim \mathcal{U}$ (-4,0) | Vague | -1.89 | -1.71 | [illegible] |
| $k_S$ | Slope of survival stress function | 1/($\mu$g.L$^{-1}$)day$^{-1}$ | $\log_{10}(k_S) \sim \mathcal{U}$ (-4,4) | Vague | -1.80 | -1.54 | [illegible] |
| $NEC$ | No Effect Concentration on daphnid survival | $\mu$g.L$^{-1}$ | $\log_{10}(NEC) \sim \mathcal{U}$ (-2,1) | Vague | 0.47 | 0.65 | [illegible] |
| $L(t, C_j)$ | Daphnid size at time $t$ and concentration $C_j$ | mm | | | | | |
| $L_0$ | Daphnid size at birth | mm | $\mathcal{N}$ (1,0.10) | [7] | 0.600 | 0.602 | [illegible] |
| $L_\infty$ | Maximum daphnid size | mm | $\mathcal{N}$ (4.8,0.59) | [8] | 3.85 | 3.92 | [illegible] |
| $k_0$ | Daphnid growth rate in the control | day$^{-1}$ | $\mathcal{N}$ (0.11,0.030) | [8] | 0.138 | 0.146 | [illegible] |
| $E_k$ | The concentration at which $k_0$ is reduced by 50 % | $\mu$g.L$^{-1}$ | $\log_{10}(E_k) \sim \mathcal{U}$ (-1,4) | | 1.26 | 1.68 | [illegible] |
| $b_k$ | Curvature coefficient | - | $\log_{10}(b_k) \sim \mathcal{U}$ (-3,3) | | -0.68 | -0.25 | [illegible] |
| $N_d(t, C_j)$ | Number of duckweed fronds per beaker at time $t$ and concentration $C_j$ | # of fronds per beaker | | | | | |
| $K_d$ | Carrying capacity for duckweeds at $t = 0$ | # of fronds per beaker | $\log_{10}(K_d) \sim \mathcal{U}$ (0,3) | [2] | 2.33 | 2.38 | [illegible] |
| $r_{d_0}$ | Intrinsic duckweed growth rate in the control | day$^{-1}$ | $\mathcal{U}$ (0,2) | [2] | 0.23 | 0.24 | [illegible] |
| $E_{r_d}$ | The concentration at which $r_{d_0}$ is reduced by 50 % | $\mu$g.L$^{-1}$ | $\log_{10}(E_{r_d}) \sim \mathcal{N}$ (2.44,0.2) | [2] | 1.96 | 2.30 | [illegible] |

| Symbol | Definition | Unit | Prior distribution* or value | Sources** | 2.5, 50, 97.5 % posterior quantiles | | |
|---|---|---|---|---|---|---|---|
| $b_{r_d}$ | Curvature coefficient | - | $\log_{10}(b_{r_d}) \sim \mathcal{U}$ (-3,3) | Vague | -1.05 | -0.91 | -0 |
| $\sigma_{N_1}$ | Standard deviation of $log_{10}$-number of algal cells per beaker in the water column | # of cells per beaker | $\mathcal{U}$ (0,5) | Vague | 0.17 | 0.18 | 0.[illegible] |
| $\sigma_{N_2}$ | Standard deviation of $log_{10}$-number of algal cells per beaker at the bottom | # of cells per beaker | $\mathcal{U}$ (0,5) | Vague | 0.21 | 0.25 | 0.[illegible] |
| $\sigma_L$ | Standard deviation of daphnid size | mm | $\mathcal{U}$ (0,2) | | 0.30 | 0.32 | 0.[illegible] |
| $\sigma_{N_d}$ | Standard deviation of $log_{10}$-number of duck-weeds fronds per beaker | # of fronds per beaker | $\mathcal{U}$ (0,2) | Vague | 0.095 | 0.10 | 0.[illegible] |

*Prior distribution: $\mathcal{N}$ stands for the normal law, $\mathcal{U}$ stands for the uniform law. **Sources: [1] Lamonica et al. 2016a, [2] Lamonica et al. 2016b, [3] DeMott 1982, [4] Grasman et al. 2009, [5] Preuss et al. 2009, [6] Villarroel at al. 1999, [7] Biron et al. 2012, [8] Billoir et al. 2008.

Table 1: Table S1: Summary of the experiments.

| Experiment | Species | Cadmium | Time | Replicate |
|---|---|---|---|---|
| 1 | Ps + Dm | 0 | 21 | 10 |
| 1 | Ps | 0 | 21 | 10 |
| 2 | Ps | 0 | 19 | 10 |
| 3 | Ps + Lm | 0 | 21 | 15 |
| 3 | Lm | 0 | 21 | 15 |
| 3 | Ps | 0 | 21 | 3 |
| 4 | Ps + Lm | 0; 11.1; 20.2; 35.5; 51.1 | 14 | 3 |
| 4 | Ps | 0; 11.1; 20.2; 35.5; 51.1 | 14 | 3 |
| 4 | Lm | 0 | 14 | 3 |
| 5 | Ps + Lm + Dm | 0; 2.25; 4.50; 6.88; 9.08 | 21 | 3 |
| 5 | Ps + Lm | 0; 2.25; 4.50; 6.88; 9.08 | 21 | 3 |
| 5 | Lm | 0; 2.25; 4.50; 6.88; 9.08 | 21 | 3 |
| 6 | Ps | 0; 26.2; 36.8; 40.8; 43.6 | 14 | 3 |

Table 3: Table S3: Data and stochastic links with variables.

| Symbol | Definition | Stochastic links with variable |
|---|---|---|
| $Y_1(t_i, C_0)$ | the observed number of algal cells in the water column at time $t_i$ and concentration $C_0$ | $\log_{10} Y_1(t_i, C_0) \sim \mathcal{N}(\log_{10}(N_1(t_i, C_0)), \sigma_{N_1})$ with $N_d = 0$ and $D_s = 0$ |
| $Y_2(t_i, C_0)$ | the observed number of algal cells at the bottom of the beaker at time $t_i$ and concentration $C_0$ | $\log_{10} Y_2(t_i, C_0) \sim \mathcal{N}(\log_{10}(N_2(t_i, C_0)), \sigma_{N_2})$ with $N_d = 0$ and $D_s = 0$ |
| $YD_1(t_i, C_0)$ | the observed number of algal cells in the water column with daphnids at time $t_i$ and concentration $C_0$ | $\log_{10} YD_1(t_i, C_0) \sim \mathcal{N}(\log_{10}(N_1(t_i, C_0)), \sigma_{N_1})$ with $N_d = 0$ |
| $YD_2(t_i, C_0)$ | the observed number of algal cells at the bottom of the beaker with daphnids at time $t_i$ and concentration $C_0$ | $\log_{10} YD_2(t_i, C_0) \sim \mathcal{N}(\log_{10}(N_2(t_i, C_0)), \sigma_{N_2})$ with $N_d = 0$ |
| $YF_2(t_i, C_0)$ | the observed number of algal cells at the bottom of the beaker with duckweeds at time $t_i$ and concentration $C_0$ | $\log_{10} YF_2(t_i, C_0) \sim \mathcal{N}(\log_{10}(N_2(t_i, C_0)), \sigma_{N_2})$ with $D_s = 0$ |
| $YF_1(t_i, C_j)$ | the observed number of algal cells in the water column with duckweeds at time $t_i$ and concentration $C_j$ | $\log_{10} YF_1(t_i, C_j) \sim \mathcal{N}(\log_{10}(N_1(t_i, C_j)), \sigma_{N_1})$ with $D_s = 0$ |
| $Ymix_1(t_i, C_0)$ | the observed number of algal cells in the water column without settling at time $t_i$ and concentration $C_0$ | $\log_{10} Ymix_1(t_i, C_0) \sim \mathcal{N}(\log_{10}(N_1(t_i, C_0)), \sigma_{N_1})$ with $N_d = 0$, $N_2 = 0$, $D_s = 0$ |
| $YFD_1(t_i, C_j)$ | the observed number of algal cells in the water column with duckweeds and with daphnids at time $t_i$ and concentration $C_j$ | $\log_{10} YFD_1(t_i, C_j) \sim \mathcal{N}(\log_{10}(N_1(t_i, C_j)), \sigma_{N_1})$ |
| $F(t_i, C_j)$ | the observed number of duckweed fronds per beaker at time $t_i$ and concentration $C_j$ | $\log_{10} F(t_i, C_j) \sim \mathcal{N}(\log_{10}(N_d(t_i, C_j)), \sigma_{N_d})$ with $N_1 = 0$, $N_2 = 0$ and $D_s = 0$ |

| Symbol | Definition | Stochastic links with variable |
|---|---|---|
| $FY(t_i, C_j)$ | the observed number of duckweed fronds with algae per beaker at time $t_i$ and concentration $C_j$ | $\log_{10} FY(t_i, C_j) \sim \mathcal{N}(\log_{10}(N_d(t_i, C_j)), \sigma_{N_d})$ with $D_s = 0$ |
| $FYD(t_i, C_j)$ | the observed number of duckweed fronds with algae and with daphnids per beaker at time $t_i$ and concentration $C_j$ | $\log_{10}(FYD(t_i, C_j)) \sim \mathcal{N}(\log_{10} N_d(t_i, C_j), \sigma_{N_d})$ |
| $W(t_i, C_j)$ | the observed number of alive daphnids per beaker at time $t_i$ and concentration $C_j$ | $W(t_i, C_j) \sim \mathcal{B}\left(\frac{S(t_i,C_j)}{S(t_{i-1},C_j)}, W(t_{i-1}, C_j)\right)$ |
| $Z(t_i, C_j)$ | the observed daphnid size at time $t_i$ and concentration $C_j$ | $Z(t_i, C_j) \sim \mathcal{N}\left(L(t_i, C_j), \sigma_L\right)$ |

Index $i$ refers to the measurement times, $t(i)$ corresponds to the $i^{\text{th}}$ measurement time points. Index $j$ refers to the tested cadmium concentrations. $\mathcal{N}$ stands for the normal law and $\mathcal{B}$ stands for the binomial law.

### 2 Figures

#### 2.1 Figure S1

#### 2.2 Figure S2 (a), (b) and (c)

#### 2.3 Figure S3

#### 2.4 Figure S4 (a) and (b)

#### 2.5 Figure S5

Fig 1: Overall graphical representation of the model. Data are symbolized by rectangles, state variables by large circles, intermediate variables by diamonds and parameters by little grey circles. Solid arrows indicate deterministic links, while dashed arrows indicate stochastic links. Thin arrows underline the time-dependencies due to the discretisation process. The dashed square indicates the variable time dependency. Sheets indicate the cadmium concentration hierarchy. Variables, data and links are detailed in ODD protocol and Tables S1 (variables and parameters) and S2 (data and links with variables).

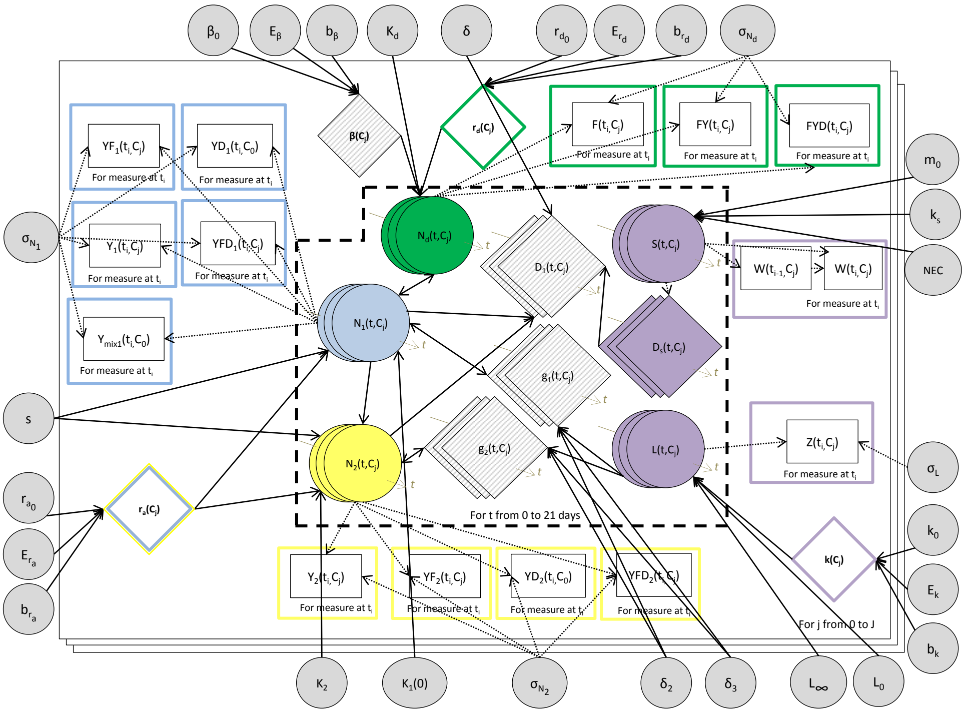


Fig 2: Representation of the fits of the model for the state variables related to duckweeds: (a) duckweed dynamics alone; (b) with algae; (c) with algae and daphnids. Symbols refer to the different tested cadmium concentrations of Experiment 5. The plain lines are the fitted model with each parameter equal to the median value of its posterior distribution at each concentrations $C_j$. The light grey area is the 95% credibility band of the predicted data in the control.

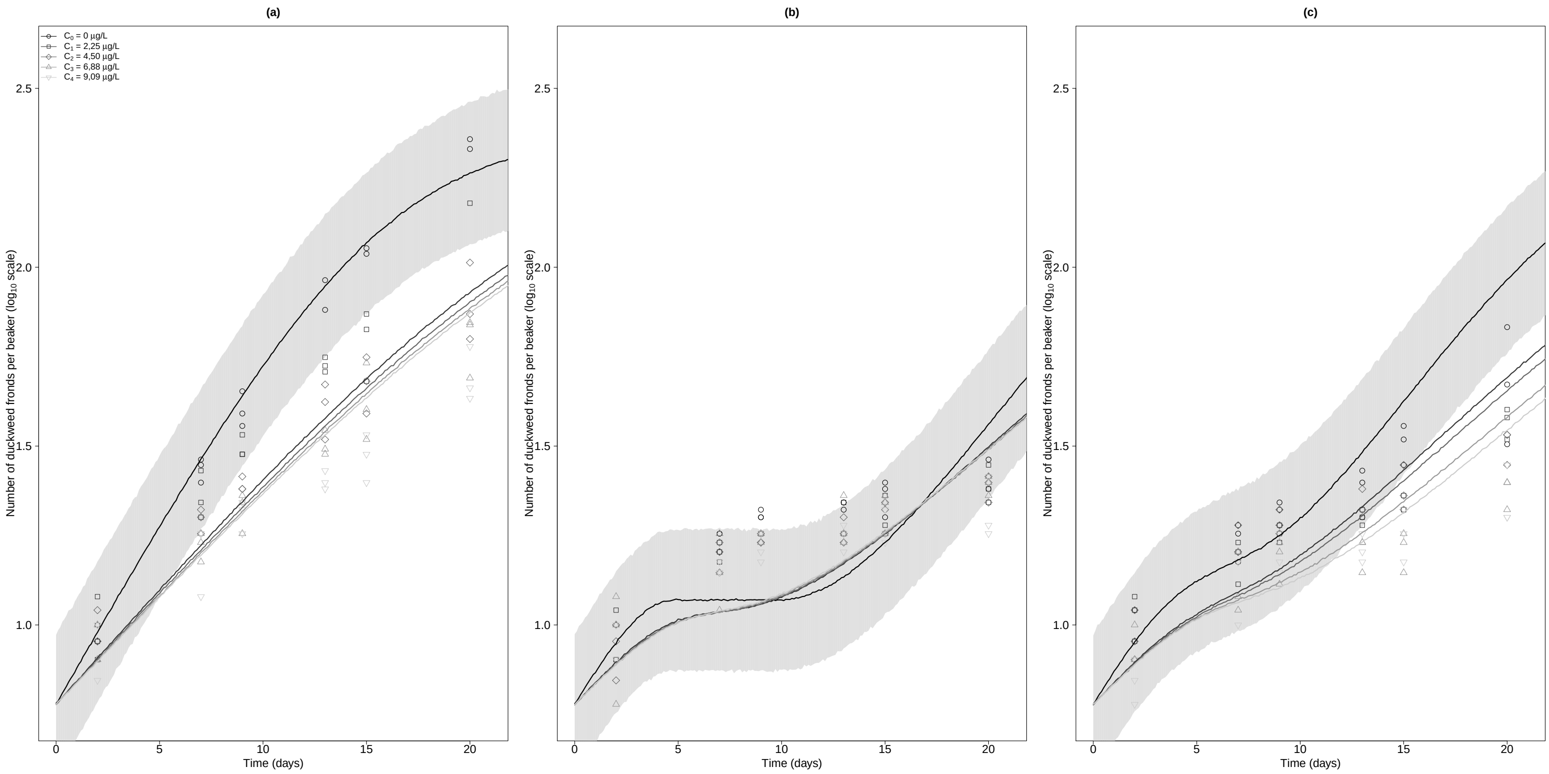

Fig 3: Fits of the model for daphnid size. Symbols refer to the different tested cadmium concentrations of Experiment 5. The plain lines are the fitted model with each parameter equal to the median value of its posterior distribution at each concentration $C_j$. The light grey area is the 95% credibility band of the predicted data in the control.

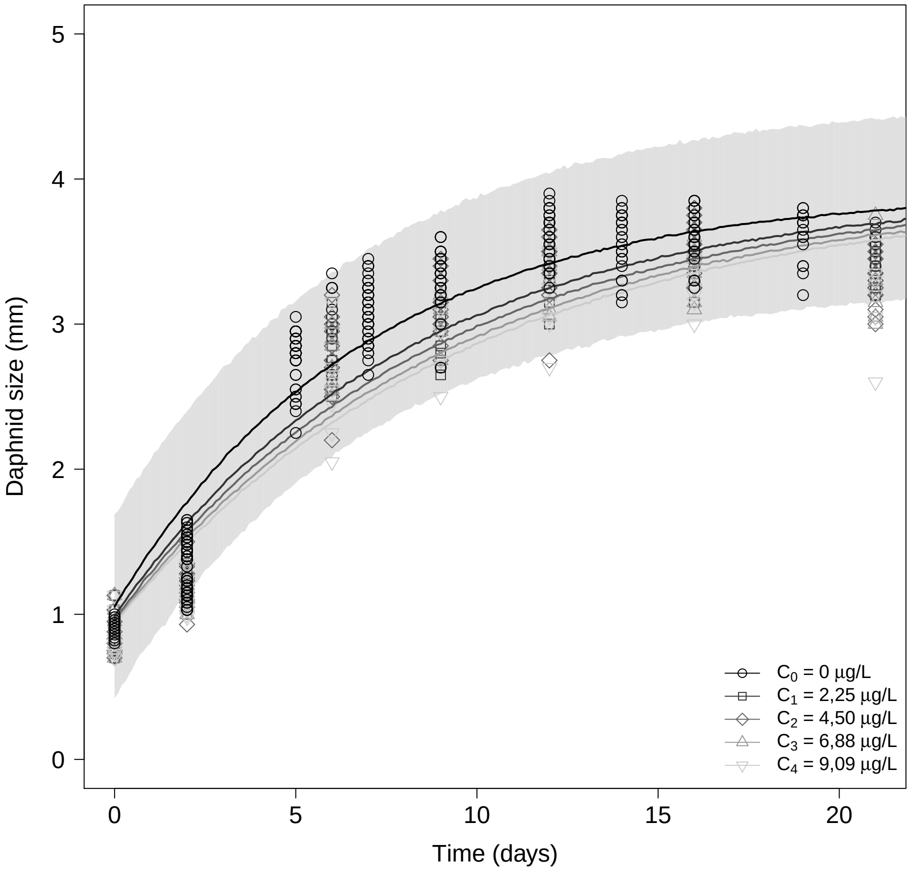

Fig 4: Comparison between prior and posterior distributions of the model parameters. The dotted lines are the prior distributions and the plain lines are the posterior distributions.

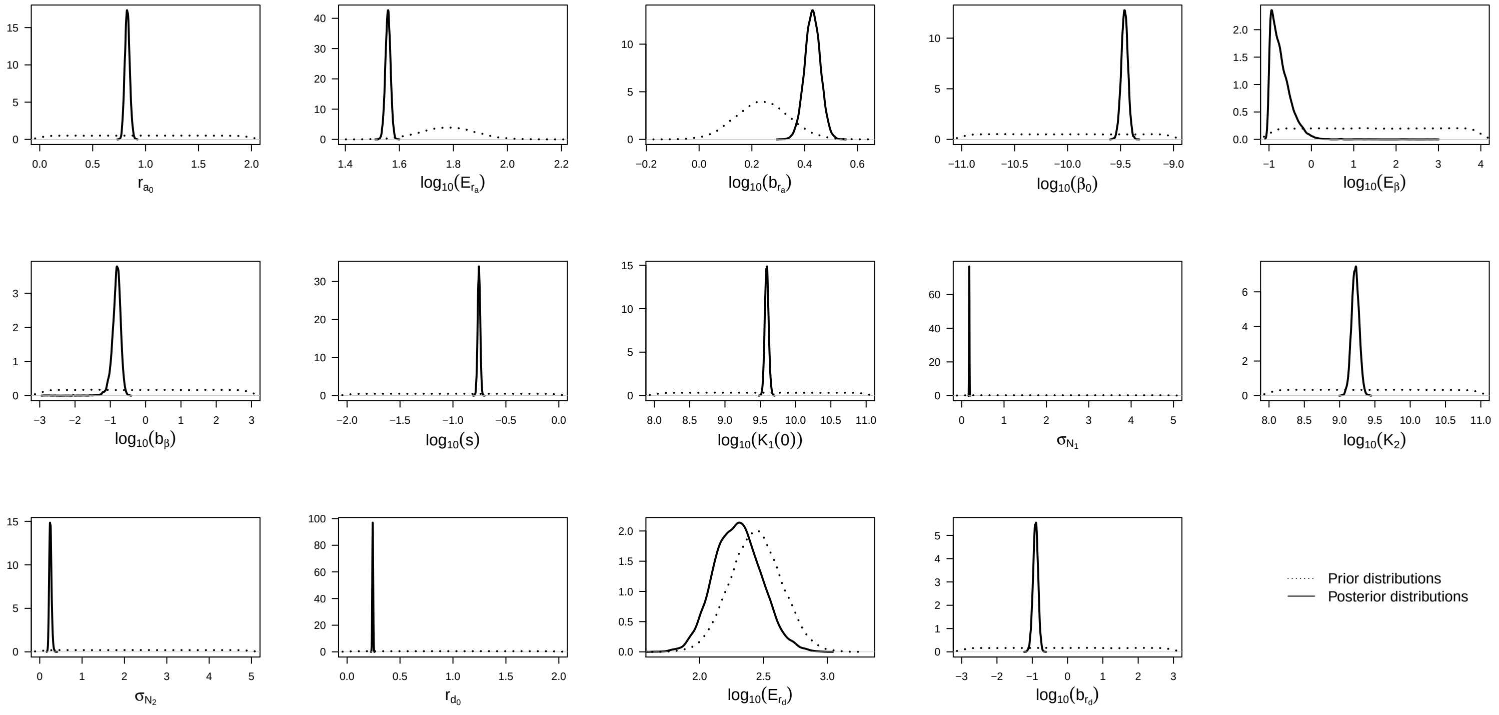


Figure S3 (b)

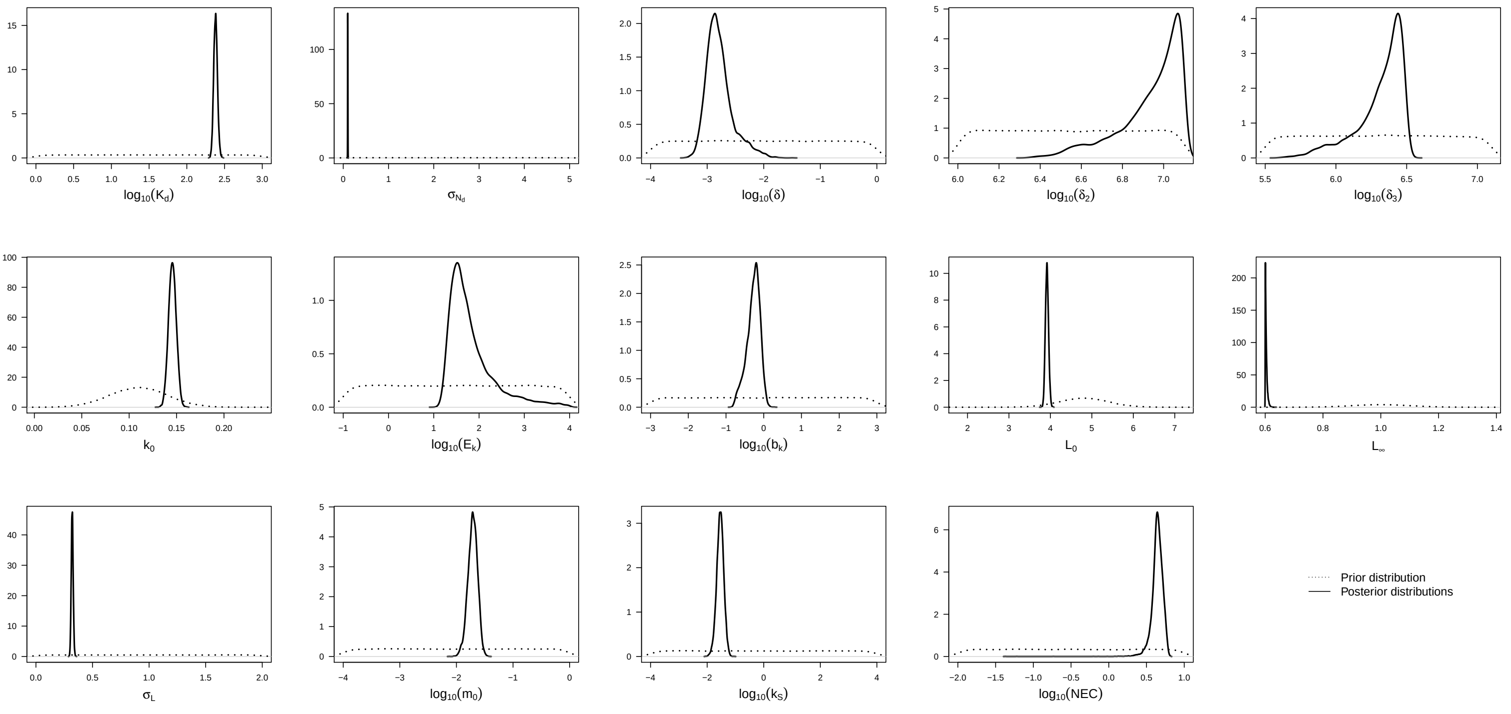

Fig 5: Boxplots of posterior distributions of the stress function parameters for each tested dataset. The reference dataset is denoted by Ref. and the partial datasets are denoted by A, B, C and D.

**Figure 3**

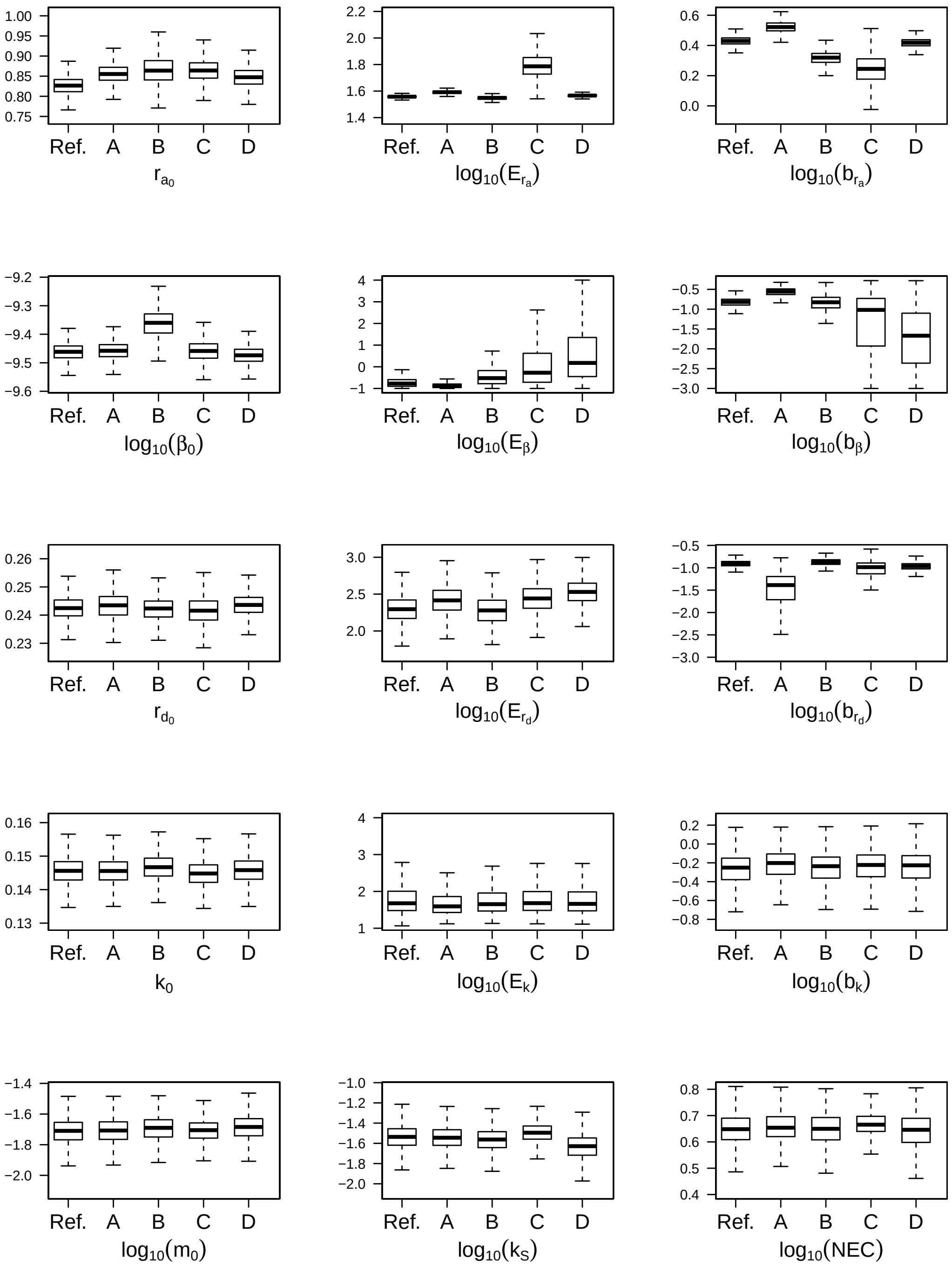

[DeMott 1982] W.R. DeMott. Feeding selectivities and relative ingestion rates of Daphnia and Bosmina. *Limnol. Oceanogr*, 27(3):518–527, 1982.

[Grasman et al. 2009] J. Grasman, E. H. van Nes, and K. Kersting. Data-directed modelling of Daphnia dynamics in a long-term micro-ecosystem experiment. *Ecological Modelling*, 220(3):343–350, February 2009.

[Lamonica et al. 2016a] D. Lamonica, U. Herbach, F. Orias, B. Clément, S. Charles, and C. Lopes. Mechanistic modelling of daphnid-algae dynamics within a laboratory microcosm. *Ecological modelling*, 320:213-230, 2016.

[Lamonica et al. 2016b] D. Lamonica, B. Clément, S. Charles, and C. Lopes. Modelling algae?duckweed interaction under chemical pressure within a laboratory microcosm. *Ecotoxicology and Environmental Safety*, 128:252-265, 2016.

[Preuss et al. 2009] T.G. Preuss, M. Hammers-Wirtz, U. Hommen, M.N. Rubach, and H.T. Ratte. Development and validation of an individual based Daphnia magna population model: The influence of crowding on population dynamics. *Ecological Modelling*, 220(3):310–329, February 2009.

[Villarroel et al. 1999] M.J. Villarroel, M.D. Ferrando, E. Sancho, and E. Andreu. Daphnia magna feeding behavior after exposure to tetradifon and recovery from intoxication. *Ecotoxicology and environmental safety*, 44(1):40–6, September 1999.
